# *In vitro* and *in vivo* antioxidant therapeutic evaluation of *Dodonaea viscosa* Jacq.

**DOI:** 10.1101/2022.04.17.488588

**Authors:** Siraj Khan, Mujeeb Ur Rehman, Muhammad Zafar Irshad Khan, Khan Muhammad, Ihsan Ul Haq, Muhammad Ijaz Khan

## Abstract

**Context:** Natural antioxidants are vital to promote health and treat critical diseased conditions in the modern healthcare system.

**Objective:** This work adds to the index of natural medicines by exploring the antioxidant potential of *Dodonaea viscosa* Jacq. (Plant-DV).

**Material and methods:** The aqueous extract of leaves and seeds from plant-DV in freshly prepared phosphate buffer is evaluated for antioxidant potential. *In vitro* antioxidant potential of the nascent and oxidatively stressed extracts was analyzed through glutathione (GSH) assay, hydrogen peroxide (H_2_O_2_) scavenging effect, glutathione-*S*-transferase (GST) assay, and catalase (CAT) activity. *In vivo* therapeutic assessment is performed in Wistar Albino rats using vitamin C as a positive control. The livers and kidneys of individual animals are probed for glutathione, glutathione-S-transferase, and catalase activities.

**Results:** Seeds have GSH contents (59.61 µM) and leaves (32.87 µM) in the fresh aqueous extracts. The hydrogen peroxide scavenging effect of leaves is superior to seeds with 17.25% and 14.18% respectively after 30 min incubation. However, oxidatively stressed extracts with Ag^I^ and Hg^II^ show declining GSH and GST levels. The plant extracts are non-toxic in rats at 5000 mg/Kg body weight. Liver and kidneys homogenate reveal an increase in GSH, GST, and CAT levels after treatment with 150 ± 2 mg/kg and 300 ± 2 mg/kg body weight plant extract compared with normal saline-treated negative and vitamin C treated positive control.

**Discussion and Conclusion:** The crude aqueous extracts of leaves and seeds of plant DV show promising antioxidant potential both in *in vitro* and *in vivo* evaluation.

## Introduction

A profound biological response of specific chemical entities in natural extracts opens new horizons to novel medicines. Natural medicines are derivatives of plants, animals, microbes, marine, and other natural sources that play an important role in the modern health care system (Sidra et al., 2014). Some biological molecules exhibit important antioxidant functions adding to the immune power of the plants and animals. Antioxidants protect the cell from oxidative damage by either scavenging or repairing the damaged molecules. The intake of enough antioxidants is supposed to protect the body against certain diseases (Jayachitra and Krithiga, 2012) like neurodegeneration, aging, viral infections, cancer and many others (Kamran et al., 2020).

Evaluation of glutathione-*S*-transferases (GST) is a potential screening method to explore the antioxidant strength of the extract. Its rich level is generally thought to offer a diverse defense system to the living cells. GST is involved in the inactivation of xenobiotics and their metabolites and catalyzing the reactive species of exogenous or endogenous origin (Sheweita, 1998). Hydrogen peroxide (H_2_O_2_) is the most abundantly produced endogenous free radical and is widely been known to produce oxidative stress. The elevated level of H_2_O_2_ is associated with suboptimal health status and needs to be warranted in any living system. Catalase prevents the cell from oxidative damage by converting hydrogen peroxide to water and oxygen (Chelikani et al., 2004). The cellular environment requires optimum GSH stores to protect itself from damage caused by the number of oxidative species. Glutathione is a naturally occurring low molecular weight non-protein thiol molecule. The presence of the sulfhydryl (SH) group has made it uniquely important by conferring an antioxidant potential. It protects the cells from free radicals and other similar oxidizing species (Jones, 2002).

Heavy metals are widespread pollutants of greater concern as they are non-degradable and thus persistent threats of oxidative toxicities. They are included in the main category of environmental pollutants as they remain in the environment for longer periods. Their accumulation is extremely hazardous to life including plants, animals, and humans (Benavides et al., 2005). A quantitative study about the antioxidant biomarkers concerning silver and mercury salts would greatly help interpret data from situations of much more complex metal exposure. Additionally, the interaction studies would present more realistic insights into the exposure of plants to these metal mixtures in the environment. The study of the effects of Ag^I^ and Hg^II^ salts on small thiols and proteins provides a good model to establish this process of conjugation at the molecular level.

The genus *Dodonaea* (Sapindaceae) - a shrub, is geographically distributed in the temperate region of Australia, South America, Mexico, Africa, India, and Florida (Mostafa et al., 2014). The genus *Dodonaea* is among 140 genera and consists of approximately 68 species (Simpson et al., 2011). Traditionally, *Dodonaea viscosa* Jacq. is used in the treatment of various ailments such as rheumatoid arthritis, diarrheas, stomach pain, skin infections, hepatic or splenic pain, uterine cramp, and other ailments involving dermatitis, smooth muscles, hemorrhoids, and sore throat (Al-Snafi, 2017).

The *Dodonaea viscosa* Jacq. (plant DV) has antioxidant (Riaz et al., 2012), anti-inflammatory (Salinas-Sánchez et al., 2012), and cytotoxic activities (Mothana et al., 2010). A study done by researcher, reported quercetin and isorhamnetin in root bark and kaempferol in its leaf extract (Getie et al., 2003). The diterpenes and nor-diterpenes are also reported in *Dodonaea viscosa* leaves (Marvilliers et al., 2020).but detailed screening of the enzymic and non-enzymic antioxidant phytochemicals remained to be explored. This study undertakes the most vital assays including glutathione (GSH) assay, hydrogen peroxide (H_2_O_2_) scavenging effect, glutathione-*S*-transferase (GST) assay, and catalase (CAT) activity to screen the antioxidant worth/value of *Dodonaea viscosa* Jacq. (Al-Snafi, 2017). Our findings show that plant DV contain antioxidant constituents that play important role in the neutralization of free radicals generated during the oxidative stress and toxic exogenous metal stress.

## Materials and methods

### Chemicals

Glutathione (GSH), 5, 5-dithio-bis (2-nitrobenzoic acid) (DTNB), 1-chloro-2,4-dinitrobenzene (CDNB), hydrogen chloride (Kolchlight), ascorbic acid, mercury II acetate (Hg(OAc)_2_), silver nitrate (AgNO_3_).

### Equipment

UV-Vis double beam spectrophotometer, HALO BD-20 (Dynamica, Australia), centrifuge (Hermle Labortechnic Germany), hot plate 400 (England), Sonicator (Sweep Zone Technology, USA), Freezer and Rotary evaporator RE200 (Bibby Sterilin Ltd England).

## Preparation of plant material and extraction

### Collection and identification

Fresh flowers and leaves of the plant were collected during the flowering season from Quaid-i-Azam University Islamabad, Pakistan. The plant was identified in the Department of Plant Sciences, Quaid-i-Azam University, Islamabad, and a voucher specimen (PHM-499) was deposited in the herbarium of medicinal plants.

### Drying

The collected plant material was sorted for any unwanted herbs and decayed or rotten plant parts. The sorted parts were thoroughly rinsed with tap water. The leaves and flowers were collected separately and air-dried at room temperature in shade, for up to three weeks until easy crumbling. Seeds were collected from the dried flowers and the thin membrane was removed by tumbling and aeration. Appropriately dried plant materials were then pulverized to get a coarsely grounded powder. The powder was individually filled in polytene jars and tightly sealed till further use.

### Extraction

The collected powder of leaves and seeds was macerated in phosphate buffer for two hrs. The homogenate was adequately stirred and filtered through a muslin cloth. The filtrate was centrifuged at room temperature at 2000 g for 10 min to remove any undissolved material. Supernatants were collected in well-closed falcon tubes freeze-dried and stored at -20 °C till further use.

## *In vitro* assessment of antioxidant potential

### Estimation of reduced glutathione (GSH)

Reduced glutathione was determined by GSH assay (Moron et al., 1979) with slight modifications as shown in Figure 1. Powder weighing 1 g/5 mL of each portion was macerated in 5% TCA solution (phosphate buffer 0. 2M, pH 8.0) and continuously stirred with a magnetic stirrer for 120 minutes. Subsequently, the extract was centrifuged at RCF=67*g* for 10 min and the supernatant was collected. The seed supernatant was serially diluted with phosphate buffer into five different concentrations at extract to phosphate buffer ratio of 7:0, 6:1, 5:2, 4:3, and 3:4. The leaves supernatant was serially diluted with phosphate buffer into five different concentrations at extract to phosphate buffer ratio of 1:5, 1:6, 1:7, 1:8, and 1:9. Normal levels of GSH in each of the five samples were determined spectrophotometrically.

**Figure 1:**
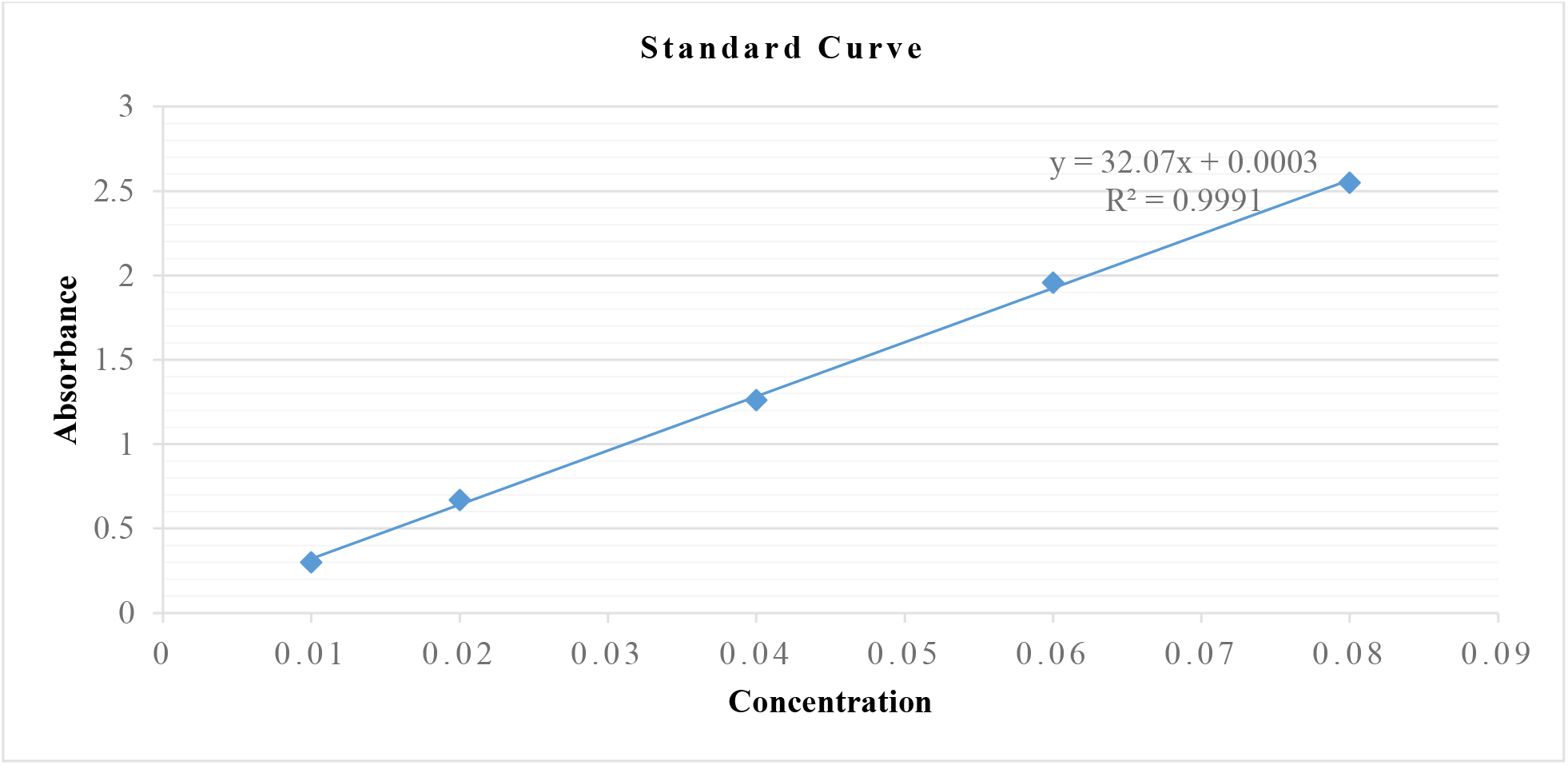
Standard curve of GSH

DTNB (2 mL) and phosphate buffer (1 mL) were used as control. The samples were prepared by mixing DTNB (0.6 mM in 0.2 M phosphate buffer, 2 mL), buffer (0.9 mL), and respective concentrations of the supernatant (0.1 mL). These mixtures were incubated for 10 min at room temperature under dark conditions. The absorbance of the assay mixture was spectrophotometrically measured at ***λ***=412 nm in triplicate, using phosphate buffer as a blank. The values were expressed as µM GSH/g of powder using a standard GSH calibration curve.

### Oxidative stress with metals

Serially diluted five different concentrations were prepared for each metal salt (mercury II acetate (Hg(OAc)_2_, silver nitrate (AgNO_3_). Stock solutions of the metal salts were prepared to be stoichiometrically equivalent to the highest concentration of the respective plant extract. The concentration of the respective plant extract was calculated from UV-vis spectrophotometric absorbance and presented in molar units. The metal salt (0.1 mL) was added to the assay mixture of different dilutions and measured spectrophotometrically after incubation of 10 min at wavelength ***λ***=412 nm, using phosphate buffer as a blank.

### Scavenging of hydrogen peroxide

The scavenging effect of extracts on H_2_O_2_ was determined over an extended period by a method reported by (Ruch et al., 1989). The leaves and seeds powder were extracted separately as per the method previously described. H_2_O_2_ scavenging effect was determined concerning different time intervals. The scavenging effect for each time interval was assessed spectrophotometrically at ***λ***=230 nm using phosphate buffer (0.1 M, pH 7.4) as a blank. The samples were analyzed and compared with the control (ascorbic acid).

Briefly,10 µL of plant supernatant was added to the H_2_O_2_ solution (40 mM; 0.6 mL) and the total volume was adjusted to 3 mL. The absorbance was recorded at ***λ***=230 nm in a spectrophotometer. The H_2_O_2_ scavenging value of the plant extracts was calculated as;

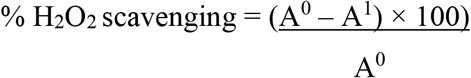

A^0^ - Absorbance of control

A^1^ - Absorbance in the presence of plant extract

### Glutathione-S-Transferase estimation

Glutathione-*S*-transferase was assessed by GST assay reported elsewhere (Habig et al., 1974). The basic principle of the assay is the ability of GST to catalyze GSH and CDNB conjugation. The activity of the enzyme was determined by observing the change in absorbance at ***λ***=340 nm. The samples were prepared by homogenizing 2 g of relevant plant-DV powder in 20 mL phosphate buffer. Homogenates were centrifuged at RCF=1680*g* for 10 min and the supernatant was collected. Different serial dilutions were prepared by adding phosphate buffer to the extract. The supernatant of seeds was serially diluted with phosphate buffer into five different concentrations at extract to phosphate buffer ratio of 7:0, 6:1, 5:2, 4:3, and 3:4. Leaves extract was serially diluted with phosphate buffer into five different dilutions at extract to phosphate buffer ratio of 1:5, 1:6, 1:7, 1:8, and 1:9. The reaction mixture contained GSH (0.1 mL), CDNB (0.1 mL) and phosphate buffer in a total volume of 2.9 mL. The reaction was initiated by the addition of enzyme extract (0.1 mL). The absorbance readings were recorded at ***λ***=340 nm in a UV-Vis spectrophotometer. The assay mixture without extract served as a control to monitor the nonspecific binding of the substrates. GST activity was calculated using the extinction co-efficient (9.6 mM^−1^cm^−1^) of the product formed and was expressed as µM of CDNB conjugated.

### Oxidative stress of extract with metals

Five different dilutions were prepared for each metal salt mercury II acetate Hg(OAc)_2_and silver nitrate AgNO_3_. Stock solutions of the metal salts were prepared to be stoichiometrically equivalent to the highest concentration of the respective plant extract. The concentration of the respective plant extract was calculated from UV-vis spectrophotometric absorbance and presented in molar units. The serial dilutions of supernatant and metals were mixed, shaken, and incubated for 10 min. After incubation, CDNB (0.1 mL), the reaction mixture (supernatant and metal, 0.1 mL), GSH (0.1 mL) and phosphate buffer (2.7 mL) were added to adjust the volume of 3 mL and recorded the absorbance at ***λ***=340 nm against phosphate buffer as a blank. The assay mixture without the extract served as a negative control to monitor the nonspecific binding of the substrates. The absorbance of positive control was also recorded. All readings were taken in triplicate. GST activity was calculated using the extinction co-efficient (9.6 mM^−1^cm^−1^) of the product formed and was expressed as µMol of CDNB conjugated/min.

### Assessment of Catalase activity

Catalase activity was determined by the catalase assay previously reported (Luck, 1974). 4 g of relevant plant DV powder were mixed in phosphate buffer (0.067 M, 20 mL, pH 7.0) through magnetic stirring. Subsequently, the extract was centrifuged at RCF=67*g* for 15 min at 4°C and the supernatant was collected. The supernatant was serially diluted with phosphate buffer into five different concentrations. Catalase activity in each diluted sample was assessed spectrophotometrically at ***λ***=240 nm. Samples were assessed against absorbance readings of the positive control. Phosphate buffer and supernatant were considered blank for each sample and served as a negative control. H_2_O_2_-phosphate buffer (2 mM) was used as a positive control. In the experimental cuvette, 40 μl enzyme extract was added to the H_2_O_2_-phosphate buffer (3 mL) and mixed thoroughly. The decline in absorbance by 0.05 units was noted at ***λ***=240 nm.

The amount of enzyme unit was calculated as the decrease in absorbance at ***λ***=240 nm by 0.05 units.

### *In vivo* assessment of the antioxidant effect

#### Animals

Wistar Albino rats (weight; 150 g and 200 g) were used in the current study to evaluate *in vivo* antioxidant activity. All experimental animals were purchased from the National Institute of Health (NIH), Islamabad Pakistan. Antioxidant activities were performed at a constant temperature of 25 ± 2°C in a pathogen-free zone of the Department of Pharmacy, Quaid-i-Azam University, Islamabad (Pakistan), as per the standard procedure for the care and use of laboratory animals (Quaid-i-Azam University). The animals were housed in plastic cages (6 rats per cage) having free contact with food and water. All animals n=48 were categorized into four groups comprising control and treated groups. Group 1 negative control was treated with normal saline, group 2 positive control with ascorbic acid 50 ± 2 mg/Kg, group 3 and group 4 were each treated with plant extracts of 150 ± 2 mg/Kg and 300 ± 2 mg/Kg respectively. Each dose of extract was dissolved in 1 ml distilled water and administered orally with the help of oral gavage. Carbon tetrachloride (CCl_4_) 0.5 ml/kg body weight 20% CCl_4_/olive oil was injected intraperitoneally into rats twice a week for 8 weeks to each animal. After 24 h from the last dose, all animals were anesthetized with chloroform and sacrificed by cervical dislocation. The liver and kidneys from each animal were collected, washed with ice-cold saline, patted dry, and weighed. Extracted tissues were individually homogenized in Tris HCl buffer (pH 7.4) to prepare homogenate. Each tissue homogenate was centrifuged at RCF=1075*g* at 4°C for 15 min. The supernatant was collected and used for the estimation of reduced glutathione (GSH), glutathione-S-transferase (GST), and catalase (CAT).

#### Acute toxicity test (LD_50_)

The acute toxicity (LD_50_) was performed as previously reported (Khalil et al., 2006). The test extract was administered at different doses (100mg/kg to 5000mg/kg) to different groups. Immediately after dosing, the animals were observed after every 4 hours for 4 days to find out mortality.

#### Determination of reduced glutathione (GSH)

Reduced glutathione was quantified by the GSH assay reported elsewhere (Ellman, 1959). The assay is based on the oxidation of GSH by 5,5-dithiobis (2-nitrobenzoic acid) (DTNB). DTNB and glutathione (GSH) react together producing 2-nitro-5-thiobenzoic acid (TNB) having a yellow color. Briefly, the sample was prepared by mixing DTNB (2.4 mL), buffer (0.5 mL), and respective dilution of the supernatant (0.1 mL). The GSH concentration was determined by measuring absorbance at 412 nm with a UV-Vis spectrophotometer using phosphate buffer as a blank.

#### Determination of Glutathione-S-transferase (GST)

The GST concentration was determined by the GST assay previously described (Doherty et al., 2010, Farombi et al., 2007). The GST enzyme is capable to conjugate GSH and CDNB. Concisely the GST level in liver and kidney tissues was estimated by mixing 0.1 mL of supernatant from respective homogenate and 0.1 mL of CDNB. Finally, 0.1 M phosphate buffer (pH 6.5) was added to make the final volume up to 3 mL. The reading was observed by UV-spectrophotometer at ***λ***=314 nm wavelength. Absorbance was measured in triplicate with a UV-Vis spectrophotometer at ***λ***=314 nm taking distilled water as a blank.

The animals were randomly divided into two groups (n=8/group). Administration of CCl_4_ (0.5 mL/kg 20% CCl4/olive oil) was done intraperitoneally (i.p.) twice a week for 8 weeks. At the same time, each animal was individually administered plant DV extract (150 ± 2 mg/kg and 300 ± 2 mg/kg b.w.) in distilled water orally, twice a week for 8 weeks. At the end of the 8^th^ week, 24 h after the last treatment, animals were sacrificed as mentioned above. Kidneys and liver were removed and washed in ice-cold normal saline and cryopreserved in liquid nitrogen and stored at -80 °C.

#### Determination of catalase activity

The method previously reported by (Luck, 1974) was used to determine catalase activity. Phosphate buffer (0.1 M, pH 7.4) was taken as a blank. The samples were analyzed against control. 40 µL of plant supernatant was added to the H_2_O_2_ solution (40 mM; 0.6 mL) and the total volume was made up to 3 mL and mixed thoroughly. The final reading was noted at 240 nm in triplicate.

## Results and discussion

### Assessment of GSH levels

#### Seeds exhibit high GSH potential then leaves in in vitro assays

The supernatant of seeds and leaf part of plant-DV were evaluated for normal GSH content without the influence of any exogenous or endogenous factor. The normal GSH concentration in seeds and leaf part of the plant-DV termed control is shown in Figure 2. GSH levels were calculated and presented in molar units obtained from the standard curve. The undiluted sample of seeds and leaves supernatant was estimated at 59.61 µM and 32.87 µM of GSH respectively. The GSH concentration was noted in the seed part to be 24.97 µM and 20.45 µM in leaves as the supernatant was diluted with aqueous phosphate buffer. After the determination of normal GSH levels, the relevant extracts were subjected to oxidative stress by exogenous silver nitrate and mercury II acetate. The strength of the Ag^I^ stock solution for the seeds and leaves was selected to be 58.34 µM and 35 µM respectively, to keep this metal solution stoichiometrically equivalent to the GSH concentration. The strength of the Hg^II^ stock solution for the seeds and leaves was selected to be 29.17 µM and 17.5 µM respectively, to keep the metallic solution stoichiometrically half to the GSH concentration in undiluted extracts. The spectroscopic evaluation of GSH concentration showed a decrease in undiluted and diluted seed and leaves supernatants. These metals (silver nitrate and mercury II acetate) showed depletion in each dilution of plant-DV. The depletion in GSH concentration in seed supernatant was decreased from 59.61 µM to 52.08 µM and 55.56 µM with Ag^I^ and Hg^II^ treated metals respectively in undiluted supernatant. The depletion in GSH concentration in leaves supernatant was also observed with Ag^I^ and Hg^II^ treated metals from 32.87 µM to 29.05 µM and 28.3 µM respectively in undiluted supernatant. The depletion in GSH concentration was also seen with all dilution in both seeds and leaves supernatant as shown in figure 2. Metals such as Ag^I^ and Hg^II^ induced stress to reduce GSH concentration by binding to the sulfhydryl group of GSH. The idea was based on the previous reports (Al-Snafi, 2017). GSH contains a thiol group having metal scavenging affinity. An earlier study showed that heavy metals toxicity increases with increasing concentration resulting in decreased GSH levels (Jozefczak et al., 2012). Additionally, the reduced GSH contents due to heavy metal treatment favor the production of reactive oxygen species (Gallego et al., 1996).

**Figure 2:**
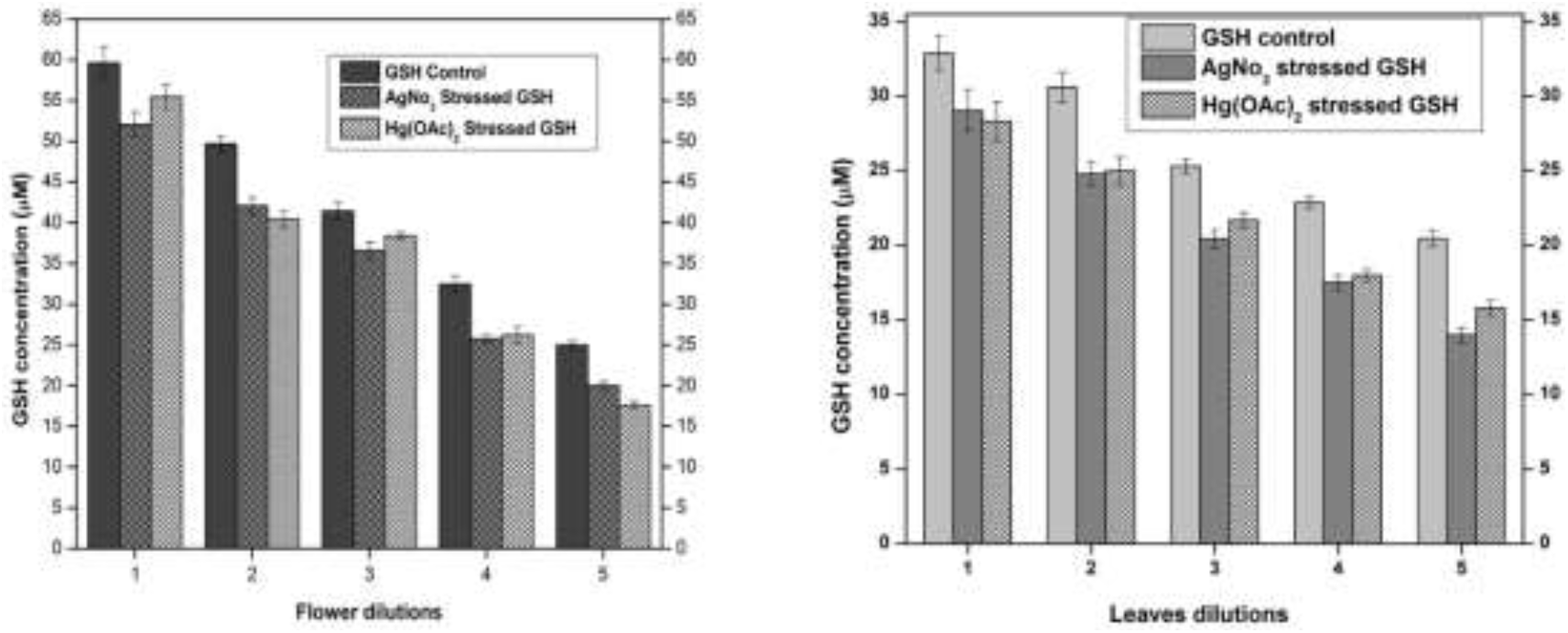
Normal and stressed Ag^I^ and Hg^II^ level of GSH expressed in µM, both flower and leaf part of plant-DV. Concentration dependent effect in five serial dilutions are determined. Values are expressed as mean ± SD (n = 3); p <0.05.

#### Seeds and leaf extract of DV-plant enhanced GSH contents in liver and kidneys of treated animals

The GSH concentration was determined in the liver and kidneys of sacrificed animals and compared with the negative control and standard as a positive control. Vitamin C was used as a standard drug for its antioxidant effect. Each group was assayed by the method previously reported by (Moron et al., 1979). As shown in Figure 3, the GSH concentration of the seed part of plant-DV was found to be 71 µM at the dose of 150 ± 2 mg/kg and 78 µM at the dose of 300 ± 2 mg/kg in the liver. However, the negative control group yielded 61 µM and the standard group/positive control treated with vitamin C showed 90 µM. In the kidney tissue, the GSH concentration of the seed extract-treated group was found to be 75 µM at the dose of 150 ± 2 mg/kg and 82 µM at the dose of 300 ± 2 mg/kg compared to the negative control group of 65 µM and vitamin C treated standard group 94 µM. The obtained results of leaves extract-treated animals indicated that the dose of 150 ± 2 mg/kg and the dose of 300 ± 2 mg/kg presented an increase in GSH concentration in liver tissue at 58 µM and 60 µM respectively compared with the negative control group 54 µM and vitamin C treated group 77 µM. In the kidney tissue, the concentration of GSH showed an increase with 63 µM at the dose of 150 ± 2 mg/kg and 69 µM at the dose of 300 ± 2 mg/kg, compared to the negative control group of 58 µM and vitamin C treated group 83 µM.

**Figure 3:**
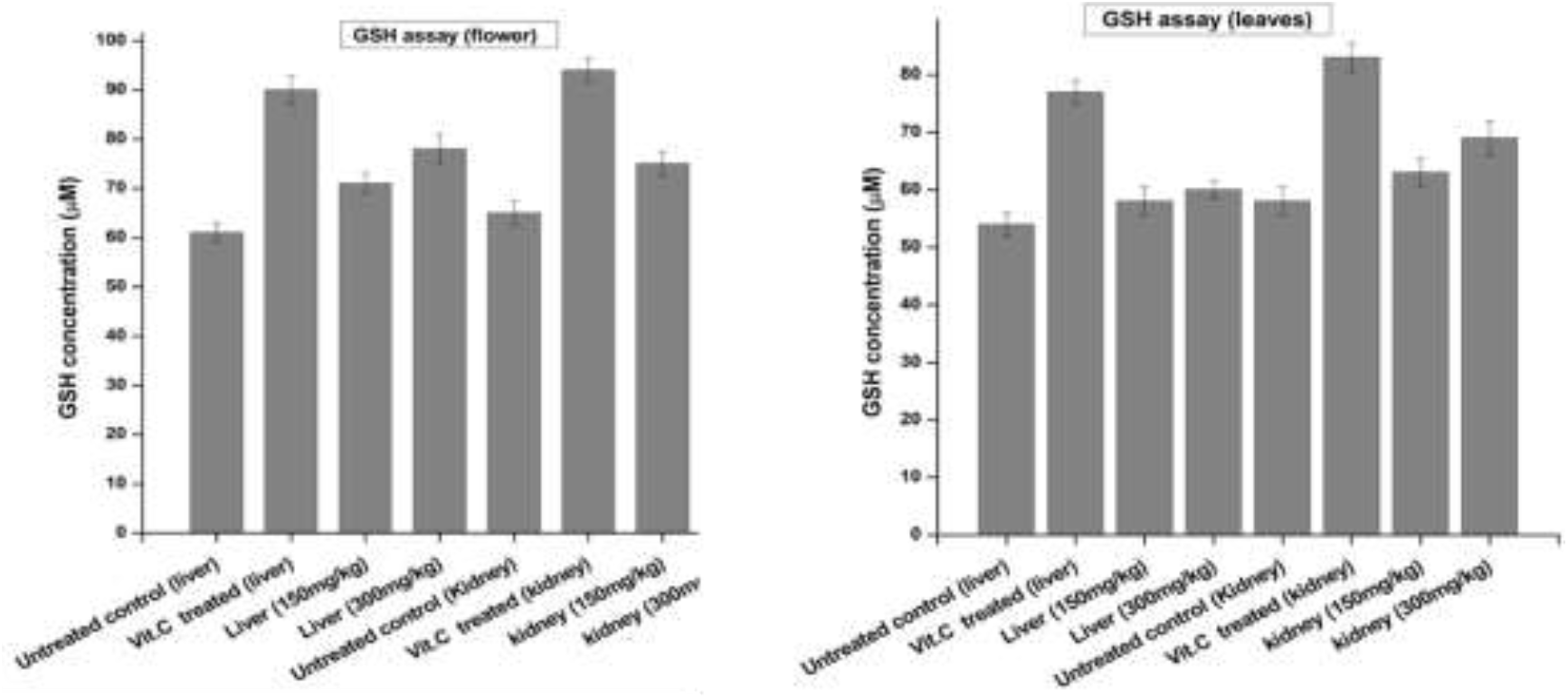
Levels of reduced glutathione (GSH) in liver and kidney of control and experimental groups of rats. Data were given as mean ± standard deviation for animals in each group; p <0.05.

Glutathione is one of the most abundant tripeptides present in the liver functioning as an enzymic antioxidant. It is primarily concerned with the removal of free radical species such as hydrogen peroxide, superoxide radicals, and alkoxy radicals. It also maintains the membrane protein thiol and works as a substrate for glutathione peroxidase and GST (Prakash et al., 2001). Our results showed that the extract caused an enhancement of GSH levels compared to the negative control group. GSH is one of the most vital antioxidant molecules exerting its antioxidant activity either through thiol conjugation or as an electron donor from its sulfhydryl group (Skenderidis et al., 2018).

### Assessment of hydrogen peroxide scavenging effect

#### The leaf part of Plant DV shows better scavenging of hydrogen peroxide than seeds

Scavenging activity of the seeds and leaf part of plant-DV were assessed for determination of the H_2_O_2_ scavenging effect. The percent scavenging effect was calculated according to the protocol reported by (Ruch et al., 1989). The percent scavenging effect was time-dependent and continuously increased as an effect of incubation time. After 1 min incubation, the scavenging effect was low in both plant parts (Figure 4). After 5 min incubation, the seeds and leaf showed a 9.7% and 12.67% scavenging effect respectively. However, after 30 min incubation, the scavenging effect of the seeds increased to 14.18% and leaves to 17.25%. Vitamin C was kept as a standard or positive control. Leaves extract showed a greater percent scavenging effect than seeds having closer results to vitamin C. The scavenging effect seems to be due to catalase, peroxidase, and the presence of flavonoids in the DV-plant (Shalaby et al., 2012).

**Figure 4:**
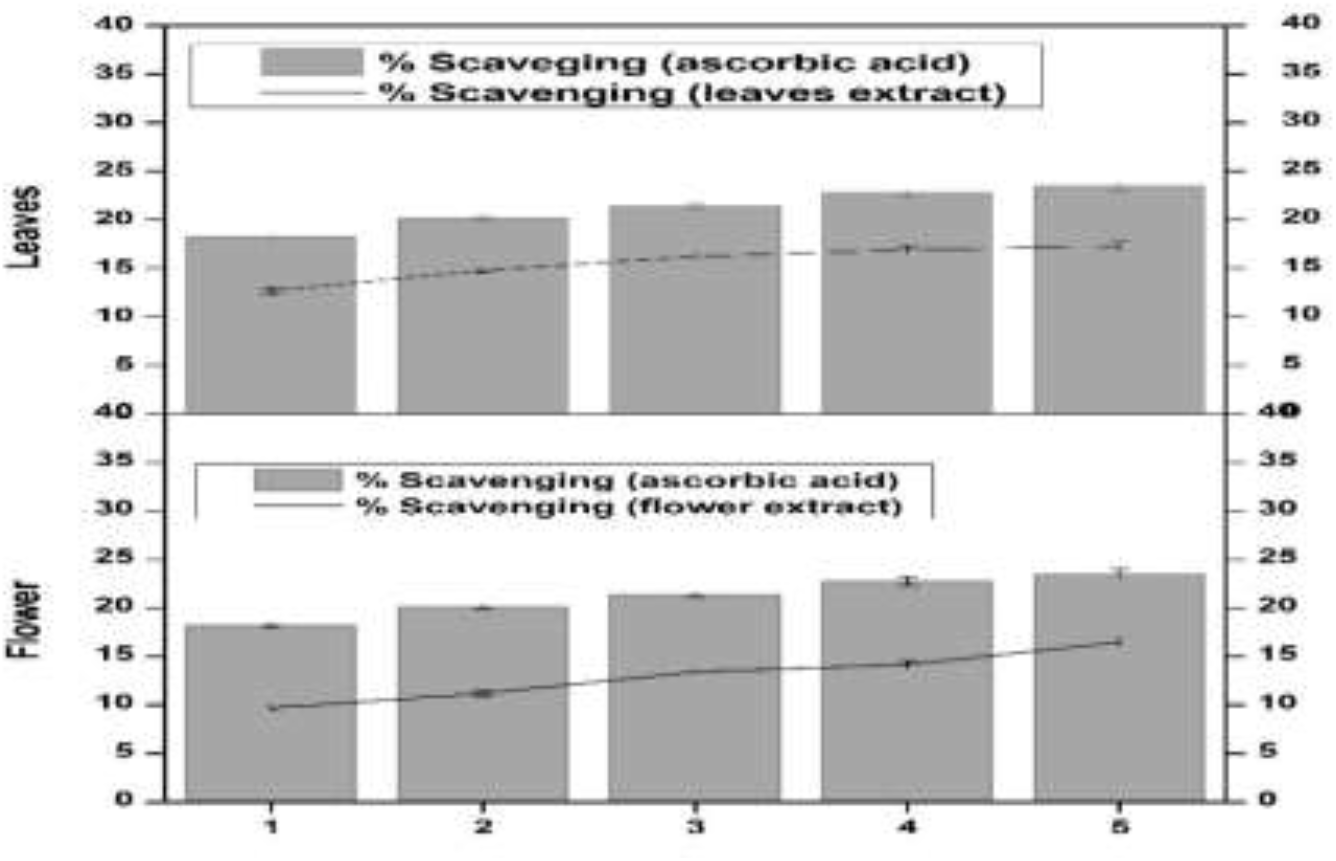
hydrogen peroxide scavenging effect of flower and leaf in plant-DV. Values are expressed as mean ± SD (n = 3); p <0.05.

### Assessment of GST levels

#### Oxidative stress reduced the GST levels in a concentration-dependent manner

The supernatant of seeds and leaves of Plant DV were evaluated for normal GST levels in serially diluted samples. GST levels were presented in molar units converted from the spectrophotometric absorbance using the extinction co-efficient (9.6 mM^−1^cm^−1^) of the product formed.

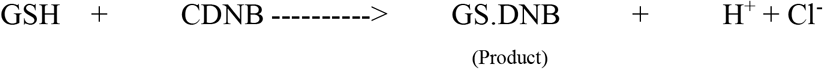

The reaction of GSH and CDNB is catalyzed by the plant enzyme GST to form the product GS-DNB having absorbance in the visible region at *λ*=340 nm. The absorbance values obtained by GS-DNB product formation are directly related to the GST potential of each supernatant dilution of the Plant DV. The extinction coefficient (9.6 mM^−1^cm^−1^) used to convert the spectrophotometric absorbance values into molar units belongs to the product GS-DNB. Normal levels of GST in each of the five dilutions were assayed spectrophotometrically by GST assay (Habig et al., 1974).

GST level in the seeds extract was found higher (153.92 µM) in the maximum supernatant concentration (DV-1) and lowest (30.41 µM) in the lowest supernatant concentration (DV-5). Similarly, the GST level in the leaf was found to be higher (157.08 µM) in the sample having maximum supernatant concentration (DV-1) and lowest (53.36 µM) in the sample having minimum supernatant concentration (DV-5) as shown in Figure 5. These findings have a positive correlation with previous studies where Plant DV has been reported to hold significant antioxidant potential (Samavati and Manoochehrizade 2013). The metal salt of Ag^I^ as an exogenous oxidizing agent was incubated with the supernatant in the same stoichiometric ratio to each serially diluted sample. GST level depletion was observed with the treatment of similarly diluted samples of Ag^I^ in the seeds of the Plant DV (Figure 5). The GST level was decreased from 153.92 µM to 85.42 µM and from 30.41 µM to 22.11 µM in the most concentrated and most diluted samples respectively. The GST level in the leaf part declined in a consistent and concentration-dependent manner. Every time with the treatment of each dilution of Ag^I^. The GST level in more concentrated samples was decreased from 157.08 µM to 85.17 µM and in diluted samples from 52.36 µM to 38.40. Treatment with Hg^II^ also showed a concentration-dependent depletion of GST contents. Every dilution of Hg^II^ depletes the GST status of the plant-DV, as shown in Figure 5. The GST level in highly concentrated samples of seeds extract was decreased from 153.92 µM to 72.92 µM and from 30.41 µM to 28.75 µM in the most diluted samples respectively. The GST level in more concentrated samples of leaf part was decreased from 157.08 µM to 96.08 µM and from 52.36 µM to 20.84 µM in diluted samples respectively.

**Figure 5:**
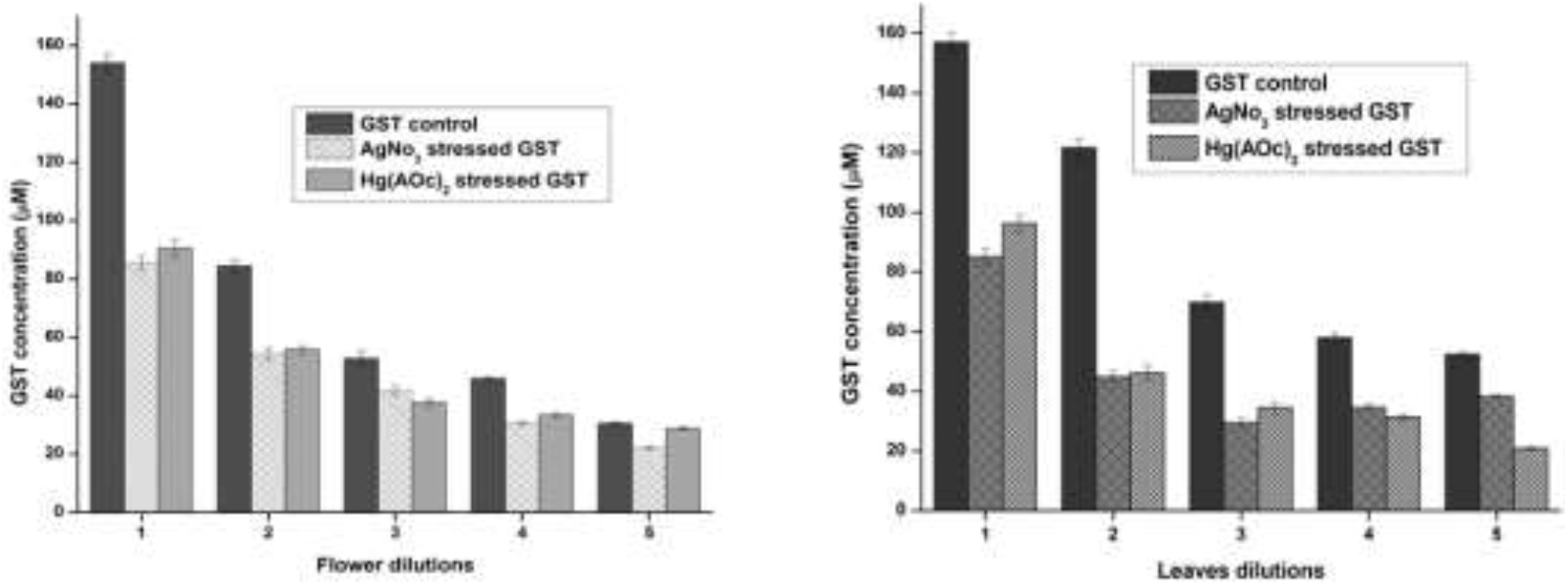
Normal and stressed Ag^I^ and Hg^II^ level of GST expressed in µM, both flower and leaf part of plant-DV. Concentration-dependent effects in five serial dilutions are determined. Values are expressed as mean ± SD (n = 3); p <0.05.

Heavy metals may inactivate enzymes by binding to a cysteine residue, thus the enzyme concentration decreases. Metal binds to sulfhydryl groups of enzymes and proteins, leading to misfolding and inhibition activity (Hossain et al., 2012). These metals might form a conjugate with GST, consequently leaving lesser GST available to catalyze the reactants i.e. GSH and CDNB.

#### Leaves showed better GST activity than seeds extract of plant-DV in in vivo evaluation

The effects of Plant DV seeds extract on *in vivo* liver and kidney GST activities are shown in Figure 6. The GST concentration in both the kidney and liver of normal animals was almost the same (220 µM). In kidneys of those animals which were administered with the 150 ± 2 mg/kg and 300 ± 2 mg/kg seeds extract, showed a significant increase in enzyme activity between the CCl_4_ treated and the extract-treated animals. The animals that were given the seed extract of 150 ± 2 mg/kg showed an increase in GST from 105 µM to 130 µM, and 300 ± 2 mg/kg showed an increase in GST from 105 µM to 164 µM. In the livers (Figure 6), the animals treated with seed extract showed a significant increase in enzyme activity between the CCl_4_ treated and the extract-treated animals. GST activity in the normal control groups was shown at 198 µM in the kidney and 219 µM in the liver. The Control group comparison showed that the normal control liver gives rise to higher enzymatic activity as compared to the normal control kidney.

**Figure 6:**
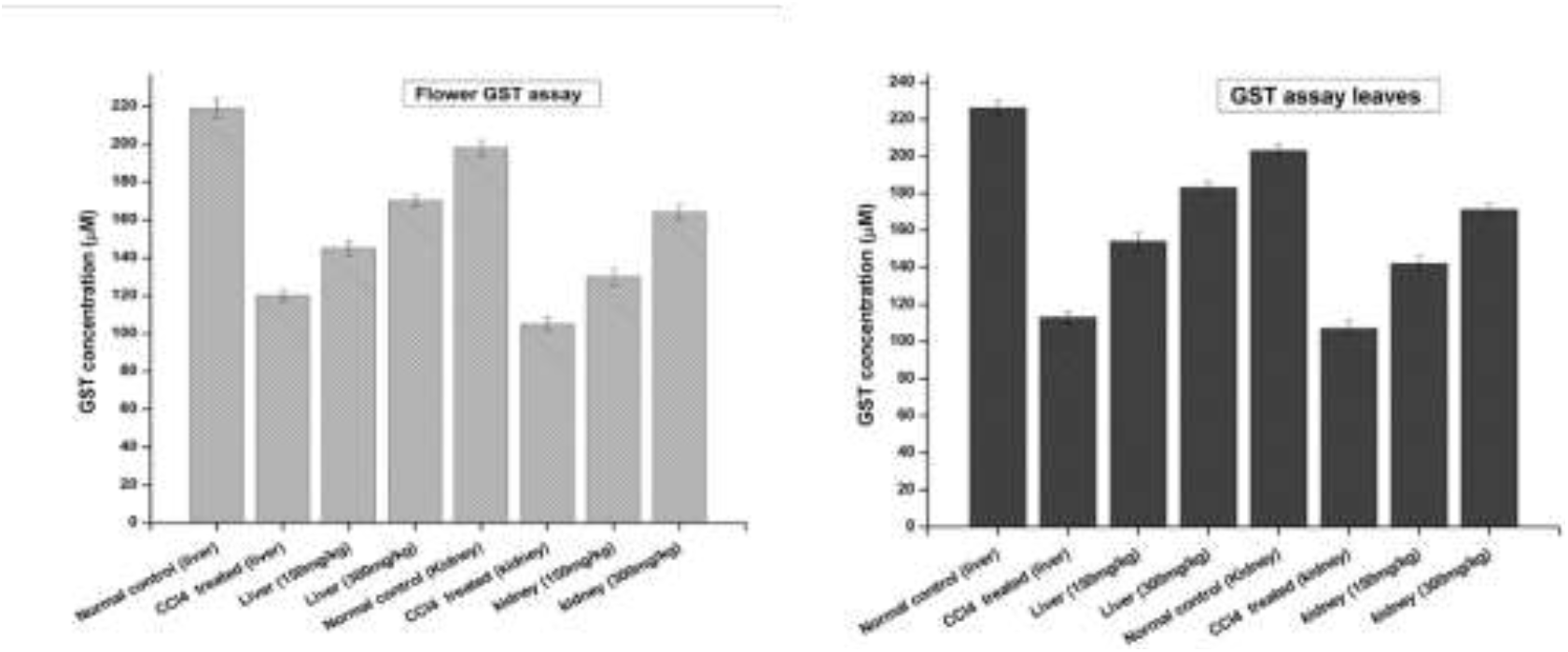
The effects of plant-DV flower and leaf extract on liver and kidney GSTs activity *in vivo* of control and experimental groups of rats. Values are expressed as mean ± SD (n = 3); p <0.05

An increase in GST activity was detected in the liver and kidney models from the animals given 150 ± 2 mg/kg and 300 ± 2 mg/kg leaves extract as shown in Figure 6. The GST activity increased by 142 µM and 171 µM for the kidney models of 150 ± 2 mg/kg and 300 ± 2 mg/kg leaves extract respectively. In the liver, as shown in Figure 6 the leaves extract-treated animals showed a substantial increase in enzyme activity at both 150 ± 2 mg/kg and 300 ± 2 mg/kg. GST is markedly shown in the liver but has also been observed in other tissues together with the kidney. Liver tissue is the major detoxification site for poisonous substances such as chemical toxins, metal ions, drugs, cancer-causing metabolites, and detoxification of endogenous noxious molecules. Liver tissue removes endogenous and exogenous toxins using GST in combination with reduced glutathione (GSH) with the electrophile center by the formation of a thioester bond among the substrate and the sulfur atom of GSH (Gulçin et al., 2018).

GST plays an important role to detoxify and metabolize many xenobiotic and endogenous compounds. Many compounds derived from plants possess antioxidant potential. *In vivo* GSTs induction is highly beneficial in protecting the cells from electrophilic insult which might be the cause of several diseases such as cancer and neurodegenerative diseases by taking electrons from macromolecules such as DNA, and proteins (van Haaften et al., 2003).

### Assessment of catalase activity

#### Leaves showed better catalase activity than seeds *in* in vitro evaluation

The supernatant of seeds and leaf part of plant-DV was assessed for the determination of catalase activity. Five serial supernatant dilutions of seeds and leaves were made by mixing the supernatant with phosphate buffer and observed for CAT activity. These dilutions spectrophotometrically provide altered absorbance in decreasing order with the decreased extract concentration. Figure 7 shows the CAT activity for all five dilutions. The amount of enzyme unit was calculated as the decrease in absorbance at a wavelength of 240 nm by 0.05 units. The activity of CAT in each of the five dilutions was evaluated by the method (Luck, 1974). As shown in Figure 7, CAT activity in the seed extract was found to be higher in the sample having maximum supernatant concentration (DV-1) as of 52.88 units/min. The CAT activity gradually declined as the supernatant was further diluted with aqueous phosphate buffer. The CAT activity in samples having minimum supernatant concentration was 9.2 units/min (DV-5). The respective CAT activity in each dilution (DV-2, DV-3, DV-4, and DV-5) is shown in Figure 7.

**Figure 7:**
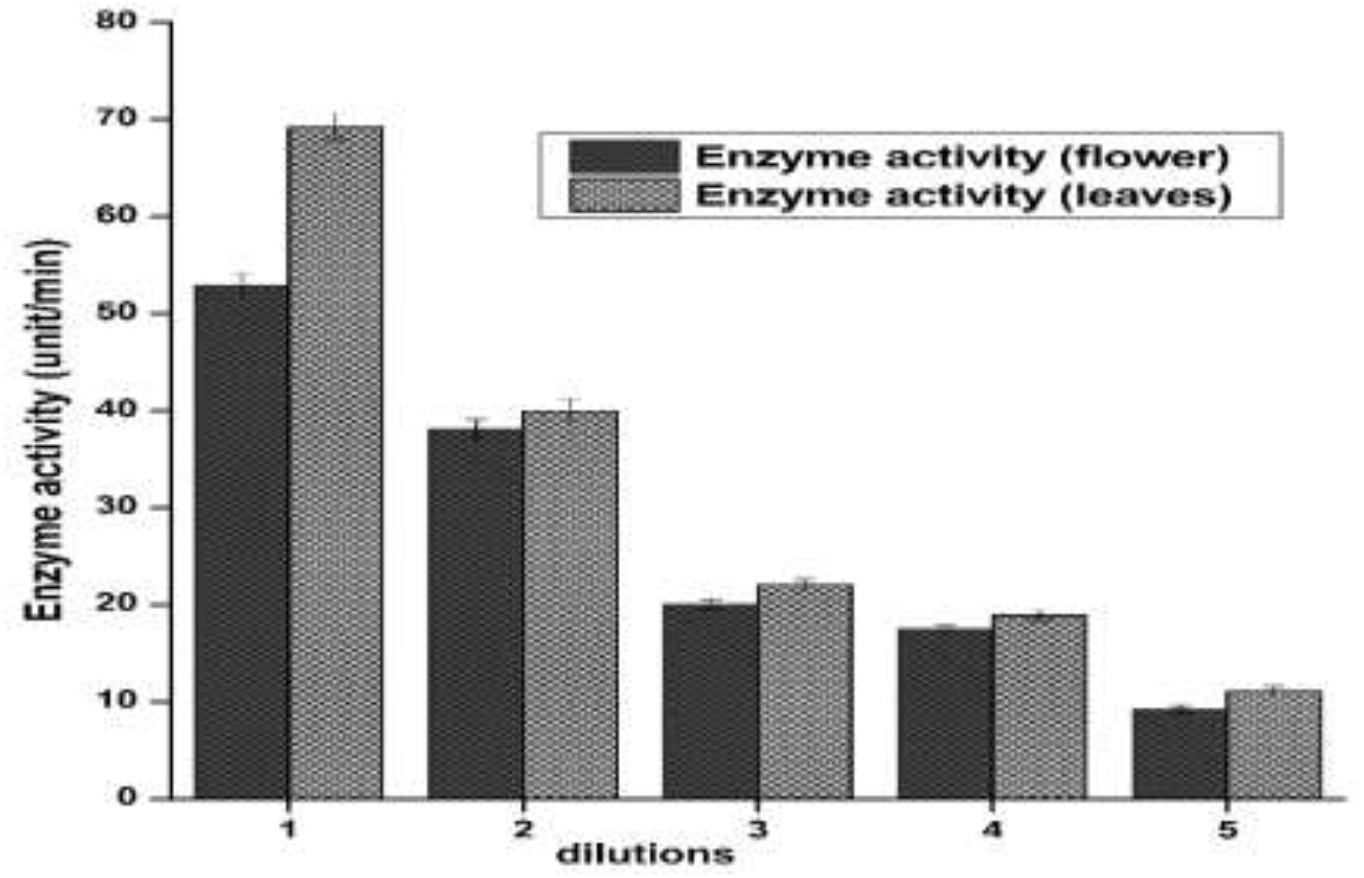
Catalase activity is expressed in unit/minute, both flower and leaf part of plant-DV. Values are expressed as mean ± SD (n = 3); p <0.05.

CAT activity in leaves extract was also found higher in the sample having maximum supernatant concentration (DV-1). The maximum supernatant concentration showed the catalytic activity of 69.2 units/min. The CAT activity gradually declined as the supernatant was further diluted with aqueous phosphate buffer. The minimum supernatant concentration denoted by DV-5 showed 11.04 unit/min catalytic activities. The respective CAT activity in each sample (DV-2, DV-3, DV-4, and DV-5) is shown the Figure 7. In more concentrated samples, there was more CAT enzyme while in a diluted sample; there may be fewer enzymes available to detoxify the free radicals. CAT catalyzed the H_2_O_2_ into H_2_O and O_2_. Due to this conversion, a decrease in absorbance was noted by the UV-Visible spectrophotometer at *λ*=240 nm. Catalase has been reported to be a common enzyme almost present in all cells that were exposed to oxygen and it decomposes hydrogen peroxide into less reactive oxygen and water (Shim et al., 2003). Previous studies have a positive correlation with these findings that plant-DV has catalase activity (MacRae and Ferguson, 1985).

#### In vivo evaluation reveals high catalase activity in leaves than in seeds

The results of catalase as shown in Figure 8 revealed an increase in liver and kidney tissue for the groups treated with seeds extract of 150 ± 2 mg/kg and 300 ± 2 mg/kg with a value of 13 unit/min, 17 unit/min and 21 unit/min, 24 unit/min, respectively compared to untreated/negative control group 28 unit/min, 32 unit/min and vitamin C treated control group 40 unit/min and 43 unit/min respectively.

**Figure 8:**
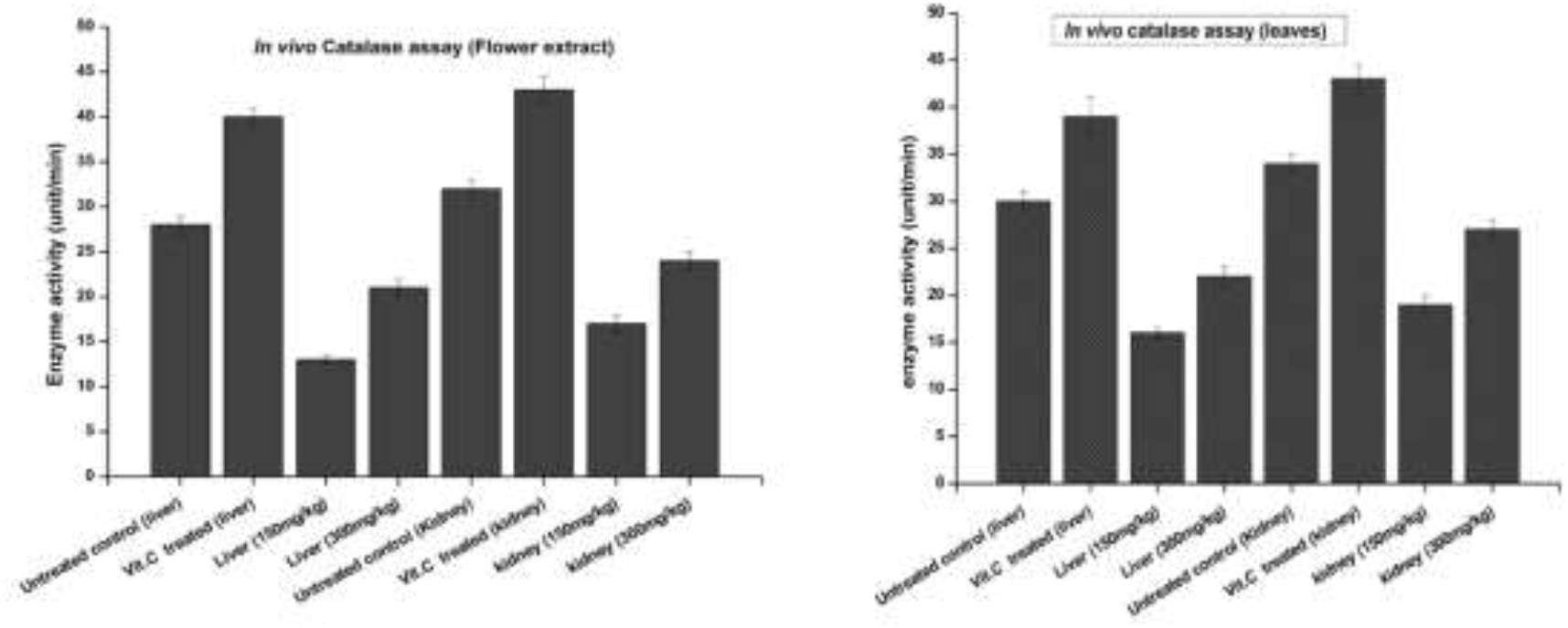
The effects of plant-DV flower and leaf extracts on the liver and kidney GSTs activity *in vivo* of control and experimental groups of rats. Values are expressed as mean ± SD (n = 3); p <0.05.

The results of catalase activity also showed an increase in liver and kidney tissue for the groups treated with leaves extract 150 ± 2 and 300 ± 2 mg/kg with the value of 16, 19 and 22, 27 unit/min, respectively compared to untreated/negative control group 30, 34 unit/min and vitamin C treated control group 39 and 43 unit/min respectively.

CAT is an enzymatic antioxidant widely distributed in all animal tissues. It decomposes hydrogen peroxide and protects the tissue from highly reactive hydroxyl radicals. CAT is part of the antioxidant enzymes that play an indispensable role in the antioxidant protective capacity of biological systems against ROS by regulating the production of free radicals and their metabolites, so the inhibition of these enzymes can cause ROS accumulation and cellular damage (Valenzuela-Cota et al., 2019). Inhibition of this enzyme may enhance sensitivity to free radical-induced cellular damage. Therefore, a reduction in the activity of CAT may lead to deleterious effects as a result of superoxide and hydrogen peroxide assimilation.

## Conclusion

This study reveals the promising antioxidant potential of leaves and seeds of Plant DV in *in vitro* and *in vivo* evaluation. *In vitro* evaluation of nascent extracts confirmed a concentration-dependent antioxidant potential of both parts having variable results in terms of various antioxidant entities and activities. Oxidatively stressed samples with Ag^I^ and Hg^II^ showed a concentration-dependent gradual decline in the antioxidant effect and efficacy of the extracts against metallic oxidation. The strong antioxidant effect of seeds and leaves extract improved the antioxidant capability of kidneys and liver of treated animals as compared to untreated and intoxicated with CCl_4_ but less potent than the standard drug, vitamin C. Bioactivity-guided evaluation of seeds and leaves from plant-DV will help to obtain pure antioxidant compounds and hence a way to novel drugs. However, the strong antioxidant potential of the crude extract can be exploited in the treatment and amelioration of various diseases to avoid the loss of micro-ingredients having fortifying, additive, or synergistic correlation.

## References

Al-Snafi, A. E. 2017. A review on Dodonaea viscosa: A potential medicinal plant. IOSR Journal of Pharmacy, 7, 10–21.

Benavides, M. P., Gallego, S. M. & Tomaro, M. L. 2005. Cadmium toxicity in plants. Brazilian journal of plant physiology, 17, 21–34.

Chelikani, P., Fita, I. & Loewen, P. C. 2004. Diversity of structures and properties among catalases. Cellular and Molecular Life Sciences CMLS, 61, 192–208.

Doherty, V., Ogunkuade, O. & Kanife, U. 2010. Biomarkers of oxidative stress and heavy metal levels as indicators of environmental pollution in some selected fishes in Lagos, Nigeria. American-Eurasian Journal of Agriculture and Environmental Science, 7, 359–365.

Ellman, G. L. 1959. Tissue sulfhydryl groups. Archives of biochemistry and biophysics, 82, 70–77.

Farombi, E. O., Adelowo, O. & Ajimoko, Y. 2007. Biomarkers of oxidative stress and heavy metal levels as indicators of environmental pollution in African cat fish (Clarias gariepinus) from Nigeria Ogun River. International journal of Environmental research and Public health, 4, 158–165.

Gallego, S. M., Benavides, M. P. & Tomaro, M. L. 1996. Effect of heavy metal ion excess on sunflower leaves: evidence for involvement of oxidative stress. Plant Science, 121, 151–159.

Getie, M., Gebre-Mariam, T., Rietz, R., Höhne, C., Huschka, C., Schmidtke, M., Abate, A. & Neubert, R. 2003. Evaluation of the anti-microbial and anti-inflammatory activities of the medicinal plants Dodonaea viscosa, Rumex nervosus and Rumex abyssinicus. Fitoterapia, 74, 139–143.

Gulçin, ă., Taslimi, P., Aygün, A., Sadeghian, N., Bastem, E., Kufrevioglu, O. I., Turkan, F. & şen, F. 2018. Antidiabetic and antiparasitic potentials: inhibition effects of some natural antioxidant compounds on α-glycosidase, α-amylase and human glutathione S-transferase enzymes. International journal of biological macromolecules, 119, 741–746.

Habig, W. H., Pabst, M. J. & Jakoby, W. B. 1974. Glutathione S-transferases the first enzymatic step in mercapturic acid formation. Journal of biological Chemistry, 249, 7130–7139.

Hossain, M. A., Piyatida, P., Da Silva, J. A. T. & Fujita, M. 2012. Molecular mechanism of heavy metal toxicity and tolerance in plants: central role of glutathione in detoxification of reactive oxygen species and methylglyoxal and in heavy metal chelation. Journal of Botany, 2012.

Jayachitra, A. & Krithiga, N. 2012. Study on antioxidant property in selected medicinal plant extract. Int. J. Med. Arom. Plants, 2, 495–500.

Jones, D. P. 2002. [11] Redox potential of GSH/GSSG couple: assay and biological significance. Methods in enzymology. Elsevier.

Jozefczak, M., Remans, T., Vangronsveld, J. & Cuypers, A. 2012. Glutathione is a key player in metal-induced oxidative stress defenses. International journal of molecular sciences, 13, 3145–3175.

Kamran, M., Kousar, R., Ullah, S., Khan, S., Umer, M. F., Rashid, H. U., Khattak, M. I. K. & Rehman, M. U. 2020. Taxonomic Distribution of Medicinal Plants for Alzheimer’s Disease: A Cue to Novel Drugs. International Journal of Alzheimer’s Disease, 2020.

Khalil, N., Sperotto, J. & Manfron, M. 2006. Antiinflammatory activity and acute toxicity of Dodonaea viscosa. Fitoterapia, 77, 478–480.

Luck, H. 1974. Estimation of catalase, methods in enzymatic analysis. Academic Press, New York.

Macrae, E. & Ferguson, I. 1985. Changes in catalase activity and hydrogen peroxide concentration in plants in response to low temperature. Physiologia Plantarum, 65, 51–56.

Marvilliers, A., Illien, B., Gros, E., Sorres, J., Kashman, Y., Thomas, H., Smadja, J. & Gauvin-Bialecki, A. 2020. Modified Clerodanes from the Essential Oil of Dodonea viscosa Leaves. Molecules, 25, 850.

Moron, M. S., Depierre, J. W. & Mannervik, B. 1979. Levels of glutathione, glutathione reductase and glutathione S-transferase activities in rat lung and liver. Biochimica et biophysica acta (BBA)-general subjects, 582, 67–78.

Mostafa, A. E., Atef, A., Mohammad, A.-E. I., Jacob, M., Cutler, S. J. & Ross, S. A. 2014. New secondary metabolites from Dodonaea viscosa. Phytochemistry Letters, 8, 10–15.

Mothana, R. A., Abdo, S. A., Hasson, S., Althawab, F., Alaghbari, S. A. & Lindequist, U. 2010. Antimicrobial, antioxidant and cytotoxic activities and phytochemical screening of some yemeni medicinal plants. Evidence-based Complementary and alternative medicine, 7, 323–330.

Prakash, J., Gupta, S., Kochupillai, V., Singh, N., Gupta, Y. & Joshi, S. 2001. Chemopreventive activity ofWithania somnifera in experimentally induced fibrosarcoma tumours in Swiss albino mice. Phytotherapy Research: An International Journal Devoted to Pharmacological and Toxicological Evaluation of Natural Product Derivatives, 15, 240–244.

Riaz, T., Abbasi, A. M., Shahzadi, T., Ajaib, M. & Khan, M. K. 2012. Phytochemical screening, free radical scavenging, antioxidant activity and phenolic content of Dodonaea viscosa. Journal of the Serbian Chemical Society, 77, 423–435.

Ruch, R. J., Cheng, S.-J. & Klaunig, J. E. 1989. Prevention of cytotoxicity and inhibition of intercellular communication by antioxidant catechins isolated from Chinese green tea. Carcinogenesis, 10, 1003–1008.

Salinas-Sánchez, D. O., Herrera-Ruiz, M., Pérez, S., Jiménez-Ferrer, E. & Zamilpa, A. 2012. Anti-inflammatory activity of hautriwaic acid isolated from Dodonaea viscosa leaves. Molecules, 17, 4292–4299.

Shalaby, N., Abd-Alla, H., Hamed, M., Al-Ghamdi, S. & Jambi, S. 2012. Flavones composition and therapeutic potential of Dodonaea viscosa against liver fibrosis. International Journal of Phytomedicine, 4, 27.

Sheweita, S. A. 1998. Heavy metal-induced changes in the Glutathione levels and Glutathione Reductase/Glutathione S-Transferase activities in the liver of male mice. International journal of toxicology, 17, 383–392.

Shim, I.-S., Momose, Y., Yamamoto, A., Kim, D.-W. & Usui, K. 2003. Inhibition of catalase activity by oxidative stress and its relationship to salicylic acid accumulation in plants. Plant Growth Regulation, 39, 285–292.

Sidra, J., Hanif, M., Khan, M. & Qadri, R. 2014. Natural products sources and their active compounds on disease prevention: a review. International Journal of Chemical and Biochemical Sciences, 6, 76–83.

Simpson, B. S., Claudie, D. J., Smith, N. M., Gerber, J. P., Mckinnon, R. A. & Semple, S. J. 2011. Flavonoids from the leaves and stems of Dodonaea polyandra: A Northern Kaanju medicinal plant. Phytochemistry, 72, 1883–1888.

Skenderidis, P., Kerasioti, E., Karkanta, E., Stagos, D., Kouretas, D., Petrotos, K., Hadjichristodoulou, C. & Tsakalof, A. 2018. Assessment of the antioxidant and antimutagenic activity of extracts from goji berry of Greek cultivation. Toxicology reports, 5, 251–257.

Valenzuela-Cota, D. F., Buitimea-Cantúa, G. V., Plascencia-Jatomea, M., Cinco-Moroyoqui, F. J., Martínez-Higuera, A. A. & Rosas-Burgos, E. C. 2019. Inhibition of the antioxidant activity of catalase and superoxide dismutase from Fusarium verticillioides exposed to a Jacquinia macrocarpa antifungal fraction. Journal of Environmental Science and Health, Part B, 54, 647–654.

Van Haaften, R. I., Haenen, G. R., Van Bladeren, P. J., Bogaards, J. J., Evelo, C. T. & Bast, A. 2003. Inhibition of various glutathione S-transferase isoenzymes by RRR-α-tocopherol. Toxicology in vitro, 17, 245–251.

